# CDK6 activity in a recurring convergent kinase network motif

**DOI:** 10.1101/2022.06.16.496486

**Authors:** Christina Gangemi, Rahkesh T Sabapathy, Harald Janovjak

## Abstract

More than 500 kinases phosphorylate ∼15% of all human proteins. The architecture of the phosphorylation network has been studied extensively, for instance to compile global interaction maps, trace signal transduction pathways or identify chemical intervention points. In contrast, systematic investigation of local motifs is rare but may provide a complementary understanding of network and kinase function. Here, we report on the occurrence, topology and experimental analysis of convergent kinase-substrate relationships (cKSRs) in which ≥two kinases phosphorylate the same substrate. Through bioinformatics analysis we found that human cKSRs are frequent and involve >80% of all kinases and >24% of all substrates. We show that cKSRs occur over a wide range of stoichiometries harnessing converging kinases that are in many instances recruited from the same family subgroup and co-expressed. Experimentally, we demonstrate at the prototypical convergent CDK4/6 kinase pair how multiple inputs at the major tumor suppressor retinoblastoma protein (RB) can hamper *in situ* analysis of individual kinases. We hypothesized that overexpression of one kinase combined with a CDK4/6 inhibitor can dissect convergence. In breast cancer cells that express high levels of CDK4, we confirmed this hypothesis and developed a high throughput compatible assay that sensitively quantifies genetically modified CDK6 variants and small molecule or protein inhibitors. Collectively, our work provides deeper insights into the phosphorylation network through the identification and analysis of kinase convergence.

## INTRODUCTION

Protein phosphorylation is the most common reversible post-translational modification in eukaryotes and an essential regulator of cell and organism physiology in health and disease. Recent experimental analysis of the phosphoproteome has revealed a large number of human protein phosphorylation sites (e.g., >200’000 sites in large mass spectrometry-based datasets, or >10’000 sites in datasets of experimentally validated kinase-substrate interactions) (Aebersold and Mann, 2016; Colinge et al., 2014; Hornbeck et al., 2015; Lemeer and Heck, 2009; Riley and Coon, 2016). The architecture of the emerging phosphorylation network has been studied extensively. Whilst many studies focused on global interaction maps or linear/interconnected signaling pathways (reviewed in (Alganem et al., 2022; Chen and Eschrich, 2014; Colinge *et al*., 2014)), less attention has been devoted to a network-wide analysis of local motifs (Invergo and Beltrao, 2018). With ∼500 kinases and ∼3’000 substrate proteins encoded in the human genome, it is expected that each kinase phosphorylates many sites and that on each substrate protein multiple sites are phosphorylated. Nodes in which multiple kinases converge on one substrate (**Fig. 1A, left**) are likely to occur in the network but their number and topologies have not been systematically investigated. These convergent network motifs are of interest not only because of their involvement in biological functions, including signal amplification, pathway crosstalk and feedback loops (**Fig. 1A, right**), but also because they can lead to challenges during experimental interrogation. For instance, quantifying the activity of a kinase *in situ* can be hampered by background phosphorylation of its substrate by additional convergent kinases. Indeed, in well-known cases of convergence, the presence of multiple inputs has resulted in limited experimental methods for functional analysis (see below).

**Figure 1:**
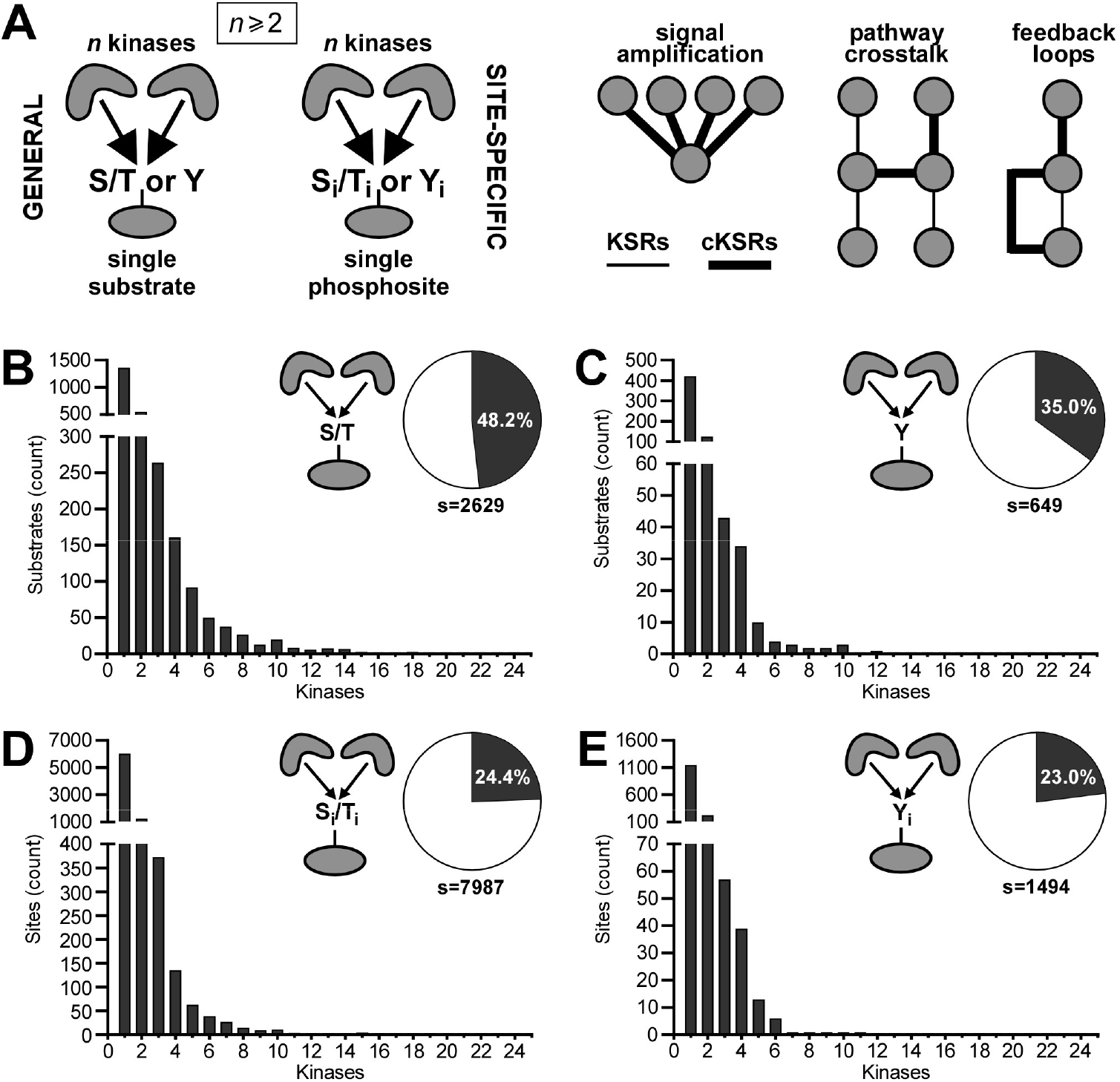
Substrate-centric analysis of convergent motifs for Ser/Thr and Tyr phosphorylation events. (**A**) cKSRs are defined as kinase-substrate interactions in which ≥two kinase phosphorylate a common substrate (general cKSRs) or a common site on a substrate (site-specific cKSRs). These interactions are involved in, e.g., amplification events, pathway crosstalk and feedback loops. (**B,C**) Distribution histograms of general cKSRs for Ser/Thr phosphorylation (B) and Tyr phosphorylation (C). Bars indicate how many substrates are phosphorylated by the indicated number of kinases. Pie charts indicate the percentage of substrates that are phosphorylated by ≥two kinases and the total number of analyzed substrates (s). (**D,E**) Distribution histograms of site-specific cKSRs for Ser/Thr phosphorylation (D) and Tyr phosphorylation (E). Bars indicate how many sites are phosphorylated by the indicated number of kinases. Pie charts indicated the percentage of sites that are phosphorylated by ≥two kinases and the total number of analyzed sites (s).

One prototypical case of phosphorylation convergence centers around the CDK4/6 serine (Ser)/threonine (Thr) kinase pair. CDK4/6 and their associated D-type cyclins (CyclinD1/D2/D3) are core components of the cell cycle machinery and regulate commitment to cycle entry (Malumbres and Barbacid, 2001; Sherr and Roberts, 1999). Unbound CDK4/6 are catalytically inactive. Assembled active CDK4/6-D-type cyclin complexes are regulated positively (e.g., by the CDK-activating complex (Fisher, 2005; Schachter et al., 2013)), negatively (e.g., by INK4 proteins (Hirai et al., 1995; Parry et al., 1999; Sherr and Roberts, 1999)), as well as positively or negatively in a context dependent manner (e.g., by KIP proteins (Blain, 2008; Cheng et al., 1999; Parry *et al*., 1999; Sherr and Roberts, 1999)). Within the cell cycle, CDK4/6 phosphorylate key regulatory sites on the major growth suppressor RB (Kaelin, 1999; Rubin, 2013). RB scavenges E2F transcription factors that are required for cell cycle progression and phosphorylation by various cyclins releases this interaction. Whilst their roles in the cell cycle and in many cases their substrate specificity are overlapping (Malumbres et al., 2004), CDK4/6 also have non-overlapping kinase activity-dependent and -independent functions outside of the cell cycle (Anders et al., 2011; Dall’Acqua et al., 2017; Hydbring et al., 2016; Tigan et al., 2016; Wang et al., 2017). Not surprisingly, CDK4/6 have received attention as major drivers of cancer and are targeted using small molecule inhibitors (Malumbres, 2014; Sherr et al., 2016). However, the absence of inhibitors that are specific for either kinase and their overlapping activity can challenge experimental analysis.

As shown previously, the presence of either CDK4 or CDK6 is sufficient to phosphorylate RB to levels that prohibit analysis of the activity of the other kinase (Malumbres *et al*., 2004; Yang et al., 2017). As a consequence, CDK4 and CDK6, in particular in the context of RB phosphorylation, are commonly functionally assayed *in vitro*, either using purified kinases and substrates, such as a recombinant RB fragment (Bockstaele et al., 2006; Jinno et al., 1999; Kitagawa et al., 1996; Paternot et al., 2003), or after immunoprecipitation (Bockstaele et al., 2009; Bockstaele *et al*., 2006). More recently, fluorescent biosensors of CDKs were developed but these require substrate-specific chemically-functionalized polypeptide probes and have not been deployed in living cells (Pellerano and Morris, 2021; Soamalala et al., 2020; Van et al., 2014). In cells, exploration of potential CDK6 substrates was conducted using gel-based electrophoretic migration shift assays that do not offer high throughput and may not in all cases provide the resolution required to visualize phosphorylation events (Anders *et al*., 2011). Finally, transcriptional reporter plasmids for signals downstream of RB often exhibit low signal-to-noise ratios (Budde et al., 2005; Chen et al., 1996a; Chen et al., 1996b). Collectively, assaying the CDK4/6-RB-axis in complex cellular environments and compatible with parallelization is currently challenging. Because of its biological/biomedical significance and these experimental limitations, the CDK4/6 pair represents an interesting test case for experimental convergence analysis.

Here, we first performed a network-wide analysis of local kinase-substrate interactions with a focus on cKSRs. We quantified their frequency, topology and kinase family subgroup distribution. We hypothesized that kinase overexpression combined with the use of a CDK4/6 inhibitor provides an approach to assay an individual kinase of the pair. Using this strategy, we were able to characterize genetically modified CDK6s and their inhibitors in breast cancer cells expressing high levels of CDK4. Collectively, our work reveals fundamental properties of cKSRs and demonstrates a possible approach to dissect convergence in cellular contexts even in the absence of kinase-specific inhibitors.

## MATERIALS AND METHODS

### Key resources table

| REAGENT or RESOURCE | SOURCE | IDENTIFIER |
| --- | --- | --- |
| **Reagents** |  |  |
| Abemaciclib (LY2835219) | AdooQ Bioscience | A12989 |
| Ribociclib (LEE011) | Selleckchem | S7440 |
| MEM | Gibco | 11095080 |
| Penicillin streptomycin | Gibco | 15140122 |
| FBS | Gibco | 26140079 |
| Insulin, recombinant human zinc solution | Gibco | 12585014 |
| PBS | Gibco | 20012050 |
| Opti-MEM | Gibco | 31985070 |
| X-tremeGENE HP DNA Transfection Reagent | Roche | 6366236001 |
| Cell Lysis Buffer (10x) | Cell Signalling Technology | 9803S |
| Glycerol | Sigma-Aldrich | G5516-500ML |
| SDS | Sigma-Aldrich | L3771-500G |
| Bromophenol blue | Sigma-Aldrich | B0126-25G |
| β-mercaptoethanol | Sigma-Aldrich | M3148-25ML |
| 20x Bolt MOPS SDS Running Buffer | Invitrogen | B0001 |
| Bolt 4-12% Bis-Tris Plus Gels 15-well | Invitrogen | NW04125BOX |
| Immun-Blot PVDF Membrane | Bio-Rad | 1620177 |
| BSA | Sigma-Aldrich | A7906-50G |
| Tween 20 | Sigma-Aldrich | P1379 |
| Clarity Western ECL Substrate | Bio-Rad | 1705060 |
| **Antibodies** |  |  |
| DYKDDDDK Tag (D6W5B) Rabbit mAb (FLAG-Tag) | Cell Signaling Technology | 14793 |
| GAPDH (D16H11) XP Rabbit mAb | Cell Signaling Technology | 5174 |
| HA-Tag (C29F4) Rabbit mAb | Cell Signaling Technology | 3724 |
| Phospho-RB (Ser780) (D59B7) Rabbit mAb | Cell Signaling Technology | 8180 |
| RB (4H1) Mouse mAb | Cell Signaling Technology | 9309 |
| Goat Anti-Mouse IgG (H+L)-HRP Conjugate | Bio-Rad | 1721011 |
| Goat Anti-Rabbit IgG (H + L)-HRP Conjugate | Bio-Rad | 1706515 |
| **Kits** |  |  |
| AlphaLISA SureFire Ultra p-RB (Ser780) | PerkinElmer | ALSU-PRB-C |
| AlphaLISA SureFire Ultra RB Total | PerkinElmer | ALSU-TRB-A |

### KSR analysis

The KSR analysis was based on a comprehensive set of experimentally determined phosphorylation sites deposited in PhosphoSitePlus (PSP) (Hornbeck *et al*., 2015). PSP is extensive and continuously manually-curated. Phosphorylation sites were downloaded (http://www.phosphosite.org, v6.6.0.2, February 2022) and further processed to obtain cKSRs using custom macros and formulas.

The first analysis was substrate-centric. This analysis was performed for Ser/Thr and tyrosine (Tyr) phosphorylation events separately and by considering either *general* cKSRs (i.e., ≥two kinases phosphorylate the same substrate but not necessarily at the same site) or *site-specific* cKSRs (i.e., ≥two kinases phosphorylate the same substrate at the same phosphorylation site). First, PSP sites were limited to human substrate entries. Second, to eliminate duplicate pairings of kinases and substrates/sites for accurate counting, fingerprints were calculated that combined the kinase and substrate/site name (e.g., “CDK4:RB” or “CDK4:RB:S780”). In a third step, the fingerprints were used to identify unique substrates/sites. In a fourth step and for each unique substrate/site, the number of active kinases was counted and their names compiled. For general cKSRs, these unique lists are given in **Supplementary Tables S1 and S2** (Ser/Thr and Tyr phosphorylation, respectively). For site-specific cKSRs, these unique lists are given in **Supplementary Tables S3 and S4** (Ser/Thr and Tyr phosphorylation, respectively). Finally, the occurrence of each possible cKSR of K:1 topology (where K denotes the number of kinases and is between 1 and 50, and 1 denotes the substrate/site) were counted. This analysis also reported on the total number of substrates/sites involved in cKSRs.

The second analysis was kinase-centric. This analysis was performed for Ser/Thr and Tyr kinases separately. Above kinase-substrate pairings were reanalyzed to produce filtered lists of kinases. The main applied filter was to include only those kinases which phosphorylate substrates that are subject to general or site-specific cKSRs (**Supplementary Tables S5-S8**). This list was then further used to retrieve kinase-specific information, such as the number of substrates that is phosphorylated by each kinase.

### Kinase family subgroup analysis

Lists containing the kinases that phosphorylate each substrate/site were generated (as in **Supplementary Tables S1-S4**). For each kinase, subgroup information was obtained from KinHub (http://kinhub.org, column: group) after correcting, where necessary and in a small number of cases, kinase nomenclature using the nomenclature list of KinHub. For each set of kinases that act on a specific substrate/site, the occurrence of members of the canonical kinase family subgroups AGC, CAMK, CK1, CMGC, RGC, STE, TK, TKL as well as atypical and “other” (Manning et al., 2002) was counted and this information encoded in a numeric string. These strings were then analyzed to reveal the number of substrates/sites that are phosphorylated by kinases from multiple family subgroups and the number of substrates/sites that are phosphorylated by ≥two kinases from at least one family subgroup.

### Identification of reciprocal phosphorylation loops

Reciprocal phosphorylation loops between two kinases were identified in the general cKSR dataset as follows: First, lists were compiled of all kinases (labelled “A” in **Supplementary Fig. S1**) that are phosphorylated by multiple kinases, e.g. an input kinase and a kinase in the reciprocal loop. Second, for each of these kinases lists of its substrates were compiled. Third, these substrate lists were compared to lists of kinases that phosphorylate kinase A to identify kinases B_1_-B_n_. Kinase A-B pairs are then compiled in **Supplementary Table S13**. For each identified phosphorylation loop, the WebOfScience database (Clarivate Analytics) was searched with keyword combinations that include the kinases and “feedback loop” or “reciprocal”. Occurrence of prior mention of the reciprocal interactions is also indicated in **Supplementary Table S13**.

### Co-expression analysis

The co-expression analysis was based on an analysis of 1393 cancer cell lines collected by the cancer cell line encyclopedia initiative (Barretina et al., 2012). Expression data of all protein-coding genes in the log_2_(TPM+1) format (where +1 denotes a pseudocount) were downloaded (http://sites.broadinstitute.org/ccle/, http://depmap.org/portal/download/, v22Q1, April 2022,) and further processed using custom macros and formulas. This analysis was performed for Ser/Thr and Tyr phosphorylation events separately and by considering either general or site-specific cKSRs. First, protein names in the expression data were cropped to only contain short HGNC compliant names. Second, the TPM values were converted into a binary format (1: expressed, 0: not expressed) using specified TPM thresholds (5, 10 and 20). Third, lists containing the kinases that phosphorylate each substrate/site were generated (as in **Supplementary Tables S1-S4**). Where necessary and in a small number of cases, kinase nomenclature was corrected using the nomenclature list of KinHub (http://kinhub.org). Fourth, for each group of kinases that act on a substrate/site, the number of cell lines was counted in which ≥two convergent kinases are co-expressed. The percentage was defined as the co-expression score and is compiled in **Supplementary Tables S9-S12**.

### Reagents

Reagents utilized are listed in the Key Resources Table. Abemaciclib (LY2835219) (AdooQ Bioscience) was diluted in dimethyl sulfoxide (DMSO) to generate a 2 mM stock solution. Ribociclib (LEE011) (Selleckchem) stock solutions were prepared at 10 mM in DMSO. Inhibitors were further diluted in cell culture media before being added to cells and equivalent amounts of DMSO were added to control wells.

### Vectors and constructs

All genes were ordered as synthetic genes (gBlocks, Integrated DNA Technologies) and designed to contain an N-terminal HA-tag (YPYDVPDYA), with the exception of Cyclin D3 that contained an N-terminal FLAG-tag (DYKDDDDK). gBlocks were amplified in polymerase chain reactions using oligonucleotides with restriction site overhangs and digested with the corresponding restriction enzymes. These were NotI and BamHI for wildtype (WT) CDK6 and p18INK4a, Xhol and BamHI for Cyclin D3, NotI and BstEII for CDK6 with N-terminal pdDronpa1, and BstEII and BamHI for CDK6 with internal loop pdDronpa1 (New England Biolabs). pcDNA3.1(-) vector (Thermo Fisher Scientific) was digested with the corresponding enzymes and ligated with amplified genes using T4 DNA ligase (Promega).

Single residue mutations were introduced in WT CDK6 using overlapping oligonucleotides designed in PrimerX (https://www.bioinformatics.org/primerx/) using the QuikChange option. Circular PCRs were performed with a high-fidelity polymerase (Q5, NEB) followed by digestion with DpnI restriction enzyme (New England Biolabs).

All genes were verified using Sanger sequencing (MicroMon, Monash University). The sequences of the engineered CDK6 fusion proteins are summarized in **Supplementary Table S14**.

### Cell culture

Human breast cancer MCF-7 cells (ATCC, HTB-22) were kindly provided by Antonella Papa (Monash University) and cultured in Minimum Essential Media (MEM, Gibco) in a humidified incubator with 5% CO_2_ atmosphere at 37°C. Medium was supplemented with 10% FBS, 0.1 mg/ml human recombinant insulin, 100 U/ml penicillin and 0.1 mg/ml streptomycin (Gibco). Transfections were conducted by seeding 500,000 cells per well in 6-well plates for immunoblotting experiments or 18,000 cells in 96-well plates for AlphaLISA assays. 24 hrs after seeding, cells were transfected with plasmid DNA using X-tremeGENE HP DNA transfection reagent (Roche) as per the manufacturer’s protocol using a total vector amount of 2 µg (6-well) or 0.1 µg (96-well). A 3:1 ratio of transfection reagent to DNA; (6 µl:2 µg) for 6-well or (0.3 µl:0.1 µg) for 96-well was used. Single construct controls were supplemented with empty pcDNA3.1(-) vector and for co-expression plasmids were added in a ratio of 1:1 or 1:1:1. Mock transfections contained empty pcDNA3.1(-) vector only. After 24 hrs, the culture medium was replaced and CDK4/6 inhibitor/s added.

### Immunoblotting

After 24 hrs of inhibitor treatment, cells were transferred onto ice and washed with cold 1x phosphate-buffered saline (PBS). 500 µl of 1x cell lysis buffer (Cell Signalling Technology) supplemented with Complete EDTA-free Protease Inhibitor Cocktail (Roche) was added for cell lysis. Lysates were collected and sonicated for 10 s, followed by incubation at 4°C with constant shaking (2000 rpm) for 30 min. Samples were then centrifuged for 20 min at 4°C before adding 4x Laemmli loading buffer (40% glycerol, 240 mM Tris/HCL, pH 6.8, 8% SDS, 0.04% bromophenol blue and 5% β-mercaptoethanol) and denaturation at 95°C for 5 min. Proteins were separated using Bolt 4-12% Bis-Tris Plus Gels (Invitrogen) run at 140 V for 60 min before being transferred onto PVDF membranes (100 V for 100 min) (Bio-Rad). Membranes were blocked in 1x TBST (1x Tris-buffered saline, 0.1% Tween) containing 5% skim milk powder for >1 hr at room temperature (rotating). Membranes were then probed with indicated primary antibodies (diluted in either 5% skim milk powder or bovine serum albumin in 1x TBST) overnight whilst rotating at 4°C. Corresponding mouse or rabbit horseradish peroxidase-conjugated secondary antibodies (diluted 1:10,000 in 1x TBST) were added for 1 hr at room temperature (Bio-Rad). Chemiluminescence was detected by adding Clarity Western ECL Substrate (1:1 of luminol and peroxide solutions) in a ChemiDoc Touch Imaging System (Bio-Rad).

### Energy transfer-based assays

AlphaLISA SureFire Ultra assay kits (PerkinElmer) were used in a two-plate two-incubation assay protocol for adherent cells according to manufacturer’s instructions. Briefly, cells were washed with 100 µl of PBS per well, followed by addition of 100 µl of 1x lysis buffer and incubated for 10 min on a plate shaker at 350 rpm. Lysate was transferred to light grey 384-well untreated AlphaPlates (PerkinElmer) by adding 10 µl per well. Both kits were used to run both assays in parallel on the same lysates from each condition. 5 µl of acceptor mix was added per well and incubated for 1 hr in the dark followed by addition of 5 µl of donor mix (under low light conditions) and incubated for a further hour in the dark. Wells were then read using a PHERAstar plate reader (BMG). Each condition was assayed in triplicates per kit.

### Data analysis and statistics

Densitometry analysis was performed on immunoblots using ImageLab 6.1 software (Bio-Rad). Kinase-substrate topology analysis was performed using equations and filters in Microsoft Excel. Reciprocal interaction analysis, kinase family analysis and co-expression analysis was performed using macros written in C in Igor Pro 6.22 (Wavemetrics). Data for **Supplementary Fig. S6** was prepared in UniProt (http://www.uniprot.org, alignment of human CDK5 and CDK6) and PyMOL 2.5 (Schrödinger; protein structures: PDB-ID 1JOW for CDK6 and 1H4L for CDK5). All other data was analysed using Prism (9.0, GraphPad).

## RESULTS

### Convergent motifs are abundant in Ser/Thr and Tyr phosphorylation

We systematically analyzed convergent phosphorylation motifs. As the data source we utilized the comprehensive PSP repository of experimentally determined phosphorylation sites (Hornbeck *et al*., 2015). >21’000 phosphorylation sites are deposited in PSP across all species, with human being the most prominent substrate type (accounting for 13’916 or 64% of sites). We limited our analysis to these human phosphorylation sites and separated Ser/Thr from Tyr kinase reactions. Using a fingerprinting approach, we identified unique substrates and for each substrate compiled the phosphorylating kinases (**Supplementary Tables S1-S4**). We found that convergent motifs are common both in Ser/Thr and Tyr phosphorylation. Of the 2629 substrates of 350 Ser/Thr kinases, 1268 (48%) are phosphorylated by more than one kinase (**Fig. 1B**). Similarly, in the case of Tyr phosphorylation by 129 kinases, 422 (65%) of 649 substrates are phosphorylated by more than one kinase (**Fig. 1C**). We next determined cKSR topologies expressed as K:1, where K denotes the number of kinases and 1 denotes the substrate. We found that 21, 10 and 6.1% of substrates are phosphorylated by two, three or four Ser/Thr kinases, and the remaining substrates by varying kinase numbers (**Fig. 1B,C**). We also found that 19, 6.6 and 5.2% of substrates are phosphorylated by two, three or four Tyr kinases, and the remaining substrates by varying kinase numbers. Thus, cKSRs are not only generally abundant but also occur with broad topologies that are somewhat comparable between Ser/Thr and Tyr phosphorylation.

The analysis conducted so far depicted general cKSRs in which ≥two kinases phosphorylate the same substrate but not necessarily at the same site. Thus, we went on to extend the analysis to site-specific cKSRs in which the same site is phosphorylated. Also in this case, we observed common convergent motifs in that 1946 (24%) of 7987 Ser/Thr sites or 344 (23%) of 1494 Tyr sites were phosphorylated by ≥two kinases (**Fig. 1D,E**). As for the general cKSRs, a wide range of topologies was observed. We found that 15, 4.7 and 1.7% of substrates are phosphorylated by two, three or four Ser/Thr kinases, and 15, 3.8 and 2.6% of substrates phosphorylated by two, three or four Tyr kinases. Collectively, these results indicate that diverse cKSRs comprise a significant fraction of the human phosphorylation landscape as mapped in the PSP. These results serve as the fundamental dataset that allows delineating key substrates onto which many kinases converge, as well as for a specific site identifying known input kinases (**Supplementary Tables S1-S4**).

In an exemplary analysis, we queried the dataset for the presence of reciprocal phosphorylation loops in which the substrate of a kinase is able to phosphorylate that kinase (rendering latter kinase into a cKSR substrate in a reciprocal phosphorylation loop; see **Supplementary Fig. S1** for a schematic of this motif). In the general cKSR dataset, we identified interactions between 33 kinases in 31 unique reciprocal loops (**Supplementary Table S13**). The majority of these interactions (41 of 62) were Ser/Thr phosphorylation events, and included well known feedback loops, such as those upstream the MAPK/ERK pathway and also those in this pathway (e.g., between MEK1 and ERK1). However, the observation that ∼half of the identified reciprocal interactions have not been described in the literature previously (**Supplementary Table S13**), demonstrates the potential of cKSR data to identify network components with specific topologies for further analysis.

### Most kinases phosphorylate cKSR substrates

We next investigated how many kinases participate in cKSRs and whether some interaction stoichiometries are more common than others (e.g., are few kinases acting on many substrates, or are many kinases acting on few substrates?). We defined the kinases of interest as those that act on substrates that are phosphorylated by ≥two kinases. We found that this criterion applied to a large fraction of human kinases (87% or 421 of 479 kinases listed in the PSP) (**Fig. 2A-D**). More specifically, of 350 Ser/Thr kinases, 324 or 298 (92 or 85%) were identified to act on general or site-specific cKSR substrates (**Fig. 2A,C, Supplementary Tables S5 and S7**). For the 129 Tyr kinases, fractions were somewhat lower but still indicated prominence (61 or 67% were associated with general or site-specific cKSR substrates; **Fig. 2B,D, Supplementary Tables S6 and S8**). We also found that kinase stoichiometries were broadly distributed. Of the identified kinases, most phosphorylated a small number of substrates whilst a small number of kinases acted on many substrates (**Fig. 2A-D**). Collectively, these results indicate that covergent motifs involve a large number of kinases through complex interactions. They allow identifying selected kinases that drive the largest number of phosphorylation events (e.g., the major signaling regulatory Ser/Thr and Tyr kinases ERK2 and SRC; **Supplementary Tables S7 and S8**).

**Figure 2:**
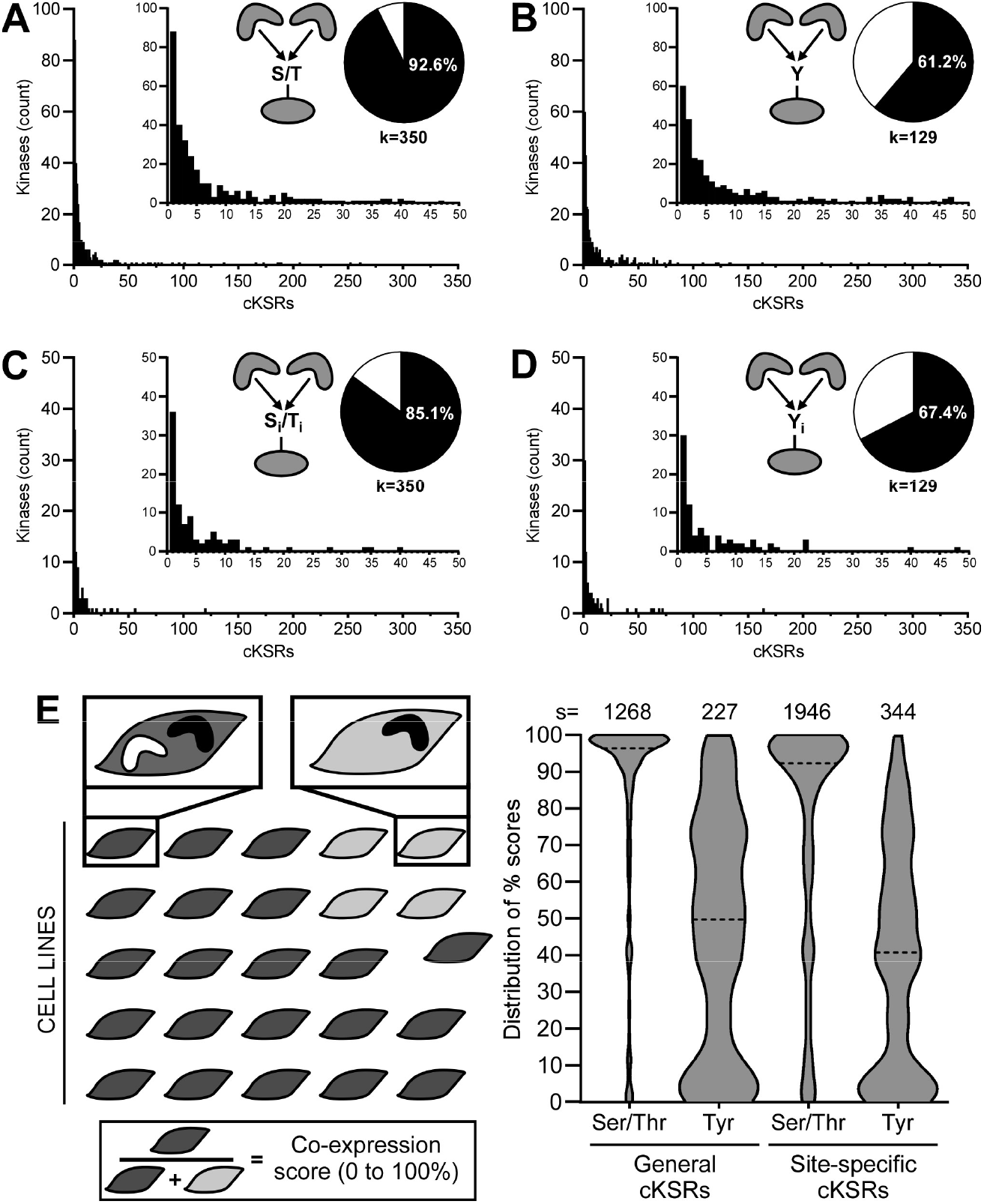
Kinase-centric analysis of convergent motifs for Ser/Thr and Tyr phosphorylation events. (**A,B**) Distribution histograms for Ser/Thr kinases (B) and Tyr kinases (C) in general cKSRs. Bars indicate how many kinases phosphorylate the indicated number of substrates. Pie charts indicate the percentage of kinases that participate in general cKSRs and the total number of kinases (k). (**C,D**) Distribution histograms for Ser/Thr kinases (B) and Tyr kinases (C) in site-specific cKSRs. Bars indicate how many kinases phosphorylate the indicated number of sites. Pie charts indicate the percentage of kinases that participate in site-specific cKSRs and the total number of kinases (k). (**E**) Co-expression analysis of converging kinases in cancer cell lines. For each group of kinases that phosphorylate the same substrate or site, the number of 1393 cancer cell lines that co-express these kinases was counted. From this, a co-expression score was calculated as the ratio of cells with (dark grey) and without (light gray) co-expression. Violin plots summarize the distribution of these scores for the analyzed number of substrates or sites (s) (dashed line: median; TPM threshold: 10).

We extended this analysis to query if kinases from multiple family subgroups converge on the same substrate or site. Our analysis focused on the major kinase family subgroups AGC, CAMK, CK1, CMGC, RGC, STE, TK, TKL as well as atypical and “other” (Manning *et al*., 2002). We found that multi-family phosphorylation is common. In the case of general cKSRs, 891 (or 70%) of 1268 Ser/Thr substrates, or 27 (or 12%) of 227 Tyr substrates were phosphorylated by kinases of ≥two families (**Supplementary Fig. S2A,B**). In the case of site-specific cKSRs, 791 (or 41%) of 1946 Ser/Thr sites, or 20 (or 5.8%) of 344 Tyr sites were phosphorylated by kinase of ≥two families (**Supplementary Fig. S2C,D**). The lower abundance of Tyr phosphorylation by multiple families needs to be interpreted with caution as it is likely due to the concentration of >90% of Tyr kinases in the TK (tyrosine kinase) group. We further investigated if multiple kinases from the same family subgroup participate in cKSRs. Strikingly, we found that this indeed applies to sizeable fractions of substrates (72.7% for Ser/Thr phosphorylation and 94.7% for Tyr phosphorylation) and sites (77.7% for Ser/Thr phosphorylation and 98% for Tyr phosphorylation) (**Supplementary Figure S3**). These observations motivated the experimental exploration of strategies to dissect the activity of closely related kinases on single substrates (see below).

Finally, we also asked if these kinases are co-expressed to relate to biological effects of cKSRs. We analyzed RNA-sequencing data from 1393 human cell lines deposited in the DepMap portal and cancer cell line encyclopedia (Barretina *et al*., 2012). Similarly to above, we compiled lists of active kinases for either each cKSR substrate or each cKSR phosphorylation site. We then examined expression levels of these kinases (TPM≥10 in **Fig. 2E**, and also TPM≥5 and 20 in **Supplementary Fig. S4**, all values are given in **Supplementary Tables S9-S12**). As a score for co-expression in each kinase group, we quantified in how many of the 1393 cell lines at least two convergent kinases are expressed (**Fig. 2E, left**). We found that cases of co-expression are generally frequent (**Fig. 2E, right**). On average and for TPM≥10, the identified Ser/Thr kinase groups are co-expressed in >1000 (or >70%) of the analyzed cell lines. Furthermore, only a small number of groups (<4%) were co-expressed more rarely (in <2.5% of cell lines). Similarly, for Tyr kinases, on average the groups were co-expressed in >930 (or >68%) of the analyzed cell lines with <9% of the groups co-expressed more rarely. Albeit co-expression is generally more prominent for Ser/Thr compared to Tyr kinases, these data collectively indicate that not only are large numbers of kinases participants in cKSRs but also that these kinases are in many cases co-expressed. The expression data tabulated in **Supplementary Tables S9-S12** allows identifying kinases and cellular contexts with and without convergence.

### Consequences of cKSRs in a prototypical kinase pair

The above analysis points to a significant occurrence of cKSRs. This promoted us to explore the consequences of convergence and to develop a strategy to dissect it for a prototypical kinase pair of the same family with specific inhibitors unavailable. This kinase pair consists of CDK4 and CDK6, two cyclin-dependent kinases that act on the major cell cycle regulator protein RB. We selected the human breast cancer cell line (MCF-7) as the main experimental system as this cell type exhibits high CDK4 levels and low CDK6 levels (TPM_CDK4_: 170, TPM_CDK6_: 0.39) (Parry *et al*., 1999; Yang *et al*., 2017) and thus offers the possibility to supplement one kinase with the other. Untreated MCF-7 cells exhibit high levels of phosphorylated RB (pRB, Ser780) attributable to CDK4 (**Fig. 3A,B**) (Li et al., 2022; Yang *et al*., 2017). We tested whether overexpression of CDK6 can elevate pRB levels but observed only a minimal effect (∼1.1-fold) in transfected cells. We also examined CDK6 variants harboring two activating mutations, S178P (Bockstaele *et al*., 2009) or R31C (Hu et al., 2011). The S178P mutant results in the cyclin-independent activation of CDK6 and thus increased cyclin-dependent activity *in vitro* (Bockstaele *et al*., 2009), whilst the R31C mutation prevents the binding of INK4 protein inhibitors resulting in sustained kinase activity (Hu *et al*., 2011). Both activating mutations, however, did not result in higher pRB levels compared to WT CDK6 (**Fig. 3A,B**). To test the possibility that D-type cyclins are limiting effects of overexpressed CDK6 (Lin et al., 2001; Parry *et al*., 1999), we co-transfected Cyclin D3. This D-type cyclin was chosen over D1 and D2 as it is known to promote high levels of kinase activity when in complex specifically with CDK6 (Lin *et al*., 2001; Parry *et al*., 1999). Cyclin D3 overexpression resulted in minor increases in pRB levels (∼1.2-fold) (**Fig. 3C,D**). The same was observed for CDK6-S178P and -R31C co-transfected with Cyclin D3 (∼1.3- and 1.2-fold elevation, resp.) (**Fig. 3C,D**). Collectively, these results indicate that overexpression of CDK6 in cells that exhibit high background activity of the convergent CDK4 does not alter pRB levels.

**Figure 3:**
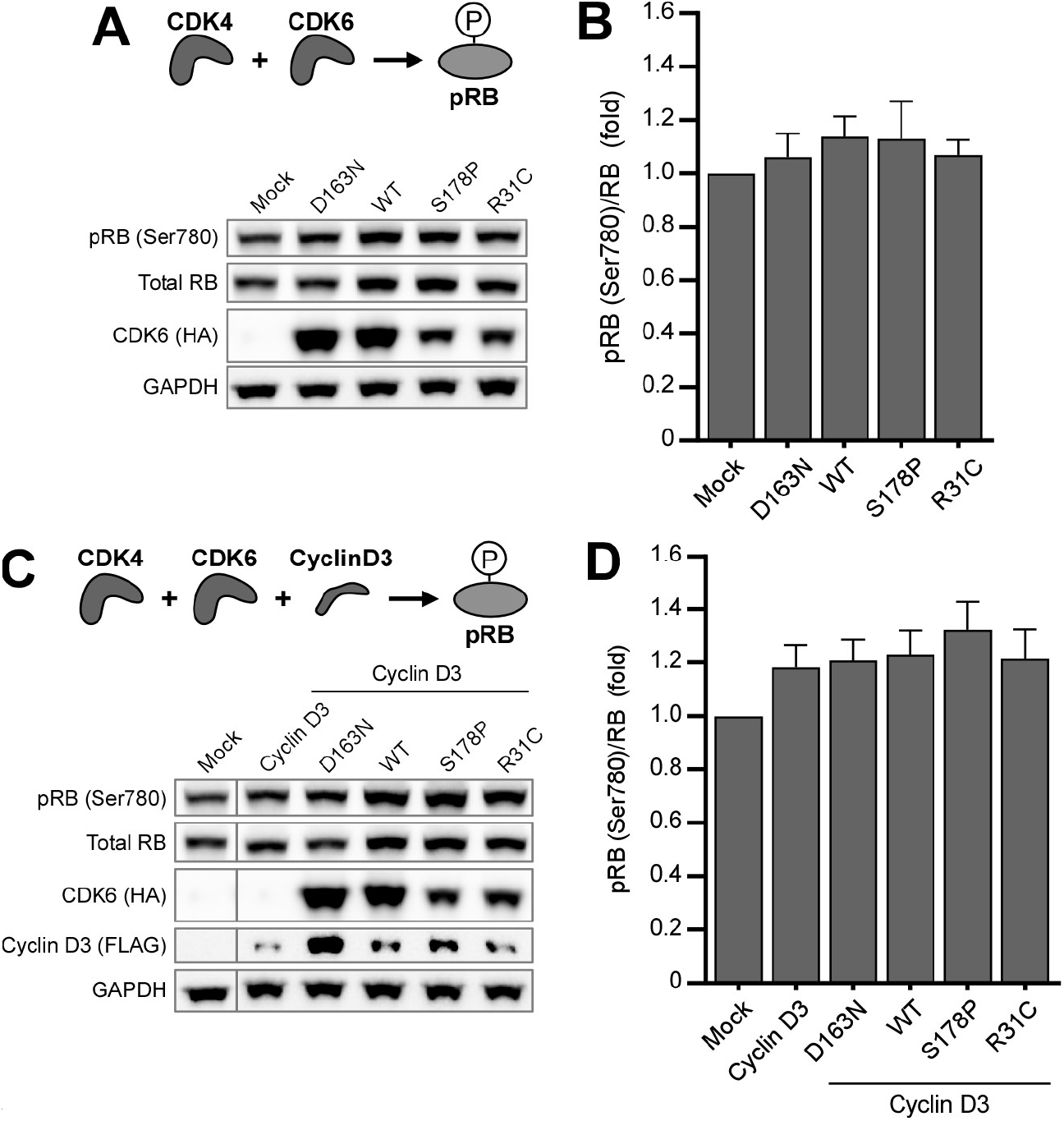
Overexpression of CDK6 and variants does not increase pRB levels. (**A**) Immunoblot analysis of pRB in MCF-7 cells expressing CDK6 and its variants. (**B**) Densitometry analysis of data shown in (A) expressed as a ratio of pRB to total RB. (**C**) Immunoblot analysis of pRB in MCF-7 cells expressing CDK6 and its variants when co-transfected with Cyclin D3. (**D**) Densitometry analysis of data shown (C) expressed as a ratio of pRB to total RB. (A and C) Representative experiments are shown. (B and D) n=3. Data are mean ± SEM.

### CDK4/6 inhibition to isolate CDK6 activity

Abemaciclib (LY2835219 [LY]) is an inhibitor of CDK4 and CDK6 with similar half maximal concentrations (IC50 2.0 and 9.9 nM) (Fassl et al., 2022; Gelbert et al., 2014). Previous studies have shown that treatment of MCF-7 cells with LY resulted in the emergence of drug resistance. This resistance was attributed to amplified CDK6 levels resulting in altered cellular inhibitor sensitivity (Cornell et al., 2019; Li *et al*., 2022; Li et al., 2018; Yang *et al*., 2017). Based on these observations, we hypothesized that application of LY may reduce background pRB levels in MCF-7 cells and that consequently overexpressed CDK6 may become functionally detectable. We first treated the cells with 0.1 µM LY as this dose was previously shown to be sufficient for inhibition of pRB signals (Yang et al, 2017). In mock-transfected cells this resulted in an almost complete reduction of RB phosphorylation (**Fig. 4A,B**). We next tested cells transfected with Cyclin D3 alone, CDK6 alone, CDK6 co-transfected with Cyclin D3, and three CDK6 variants (the loss-of-function variant D163N and above gain of function variants) also co-transfected with Cyclin D3 (**Fig. 4A,B**). Strikingly, we found that in the presence of LY overexpression of CDK6 induced pRB levels significantly (∼2.5-fold). Furthermore, co-expression of CDK6 with Cyclin D3 resulted in an even greater increase by ∼4.1-fold. Similar to our previous experiments, the S178P or R31C variants produced comparable effects to CDK6. The D163N variant as expected showed low pRB levels similar to control cells which demonstrates that effects of ectopic expression of CDK6/Cyclin D3 are kinase-dependent and attributable to active CDK6. Whilst a single dose of LY delivered the desired outcome of isolating CDK6 activity from the CDK4 background, we additionally performed dose response curves (**Fig. 4C,D**). Whilst reduction of pRB levels at increasing concentrations was observed in mock-transfected and Cyclin D3 overexpressing cells, these levels were almost entirely sustained in cells transfected with CDK6/Cyclin D3. A similar selection phenomenon was also observed using Ribociclib ([RI], LEE-011), another CDK4/6 inhibitor that is not chemically related to LY (Fassl *et al*., 2022; Zhang et al., 2014) (**Supplementary Fig. S5**). Overall, these results demonstrate that even in the absence of a specific inhibitor CDK6 activity can be isolated in a convergent motif.

**Figure 4:**
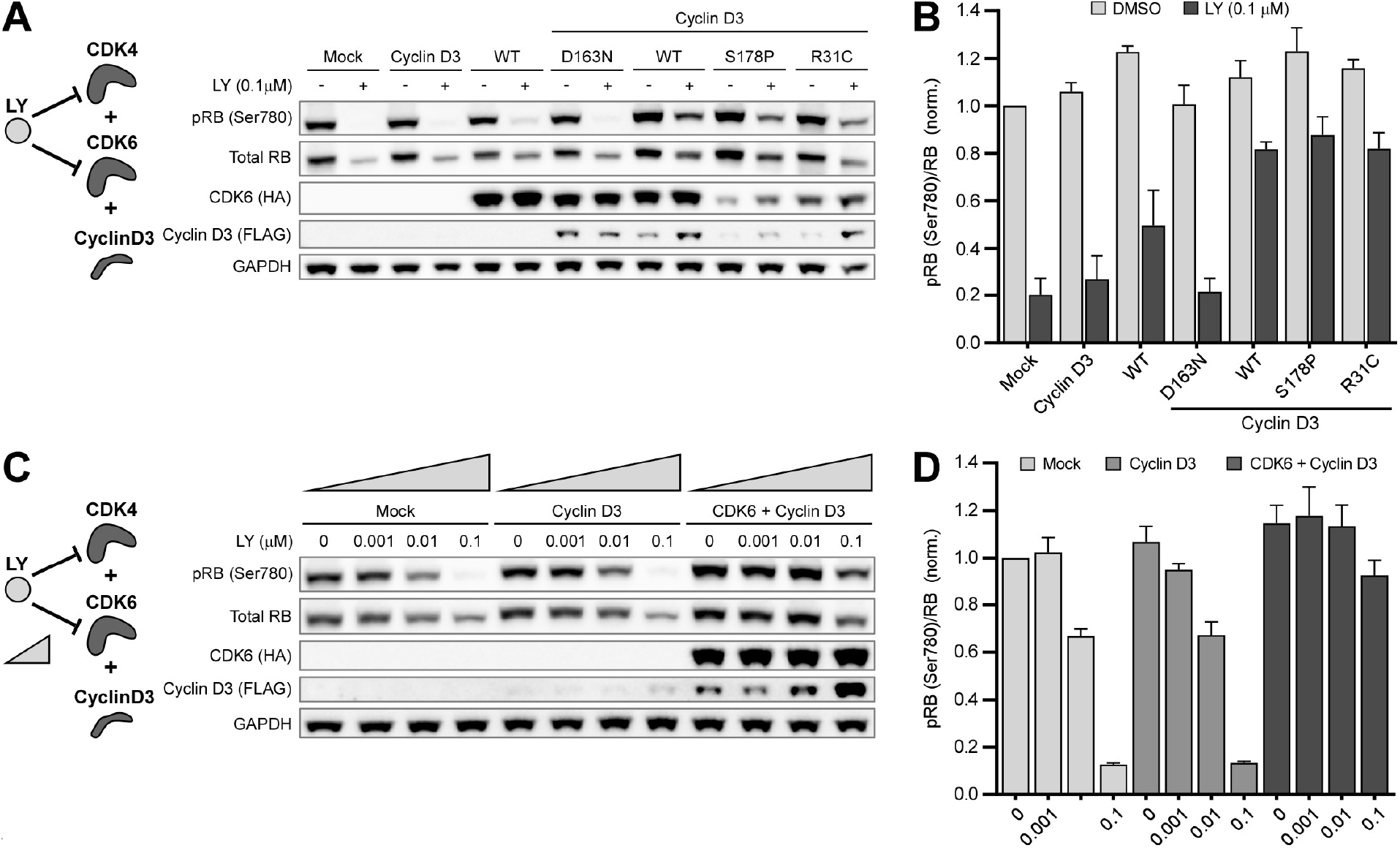
Kinase inhibition with LY reveals CDK6 activity. (**A**) Immunoblot analysis of pRB in MCF-7 cells expressing CDK6 and its variants when co-transfected with Cyclin D3, treated with LY or DMSO. (**B**) Densitometry analysis of data shown in (A) expressed as a ratio of pRB to total RB. (**C**) Immunoblot analysis of pRB in MCF-7 cells expressing CDK6 and Cyclin D3, treated with increasing concentrations of LY. (**D**) Densitometry analysis of data shown in (C) expressed as a ratio of pRB to total RB. (A and C) Representative experiments are shown. (B) n=4 and (C) n=3. Data are mean ± SEM.

### In situ assay of CDK6 variants and modulators

We have demonstrated through immunoblotting that CDK6 activity can be quantified in cells using a combination of overexpression and CDK4/6 inhibition. We went on to test if this approach can be applied in a high throughput compatible platform and in the exploration of the function of engineered CDK6 variants. We employed an energy transfer-based assay to detect pRB (Ser780) and total RB levels (**Fig. 5A**). This assay was chosen as it can be used with a greater efficiency than immunoblotting. We first tested the same conditions as above and observed consistent results. CDK6 and activating mutations displayed the greatest fold-increases in pRB levels specifically in the presence of LY (e.g., a >6-fold difference in pRB levels compared to mock inhibitor-treated cells; **Fig. 5A,B**). Also, as observed earlier, CDK6 alone only showed a minor increase and the two activating mutations overall behaved similarly.

**Figure 5:**
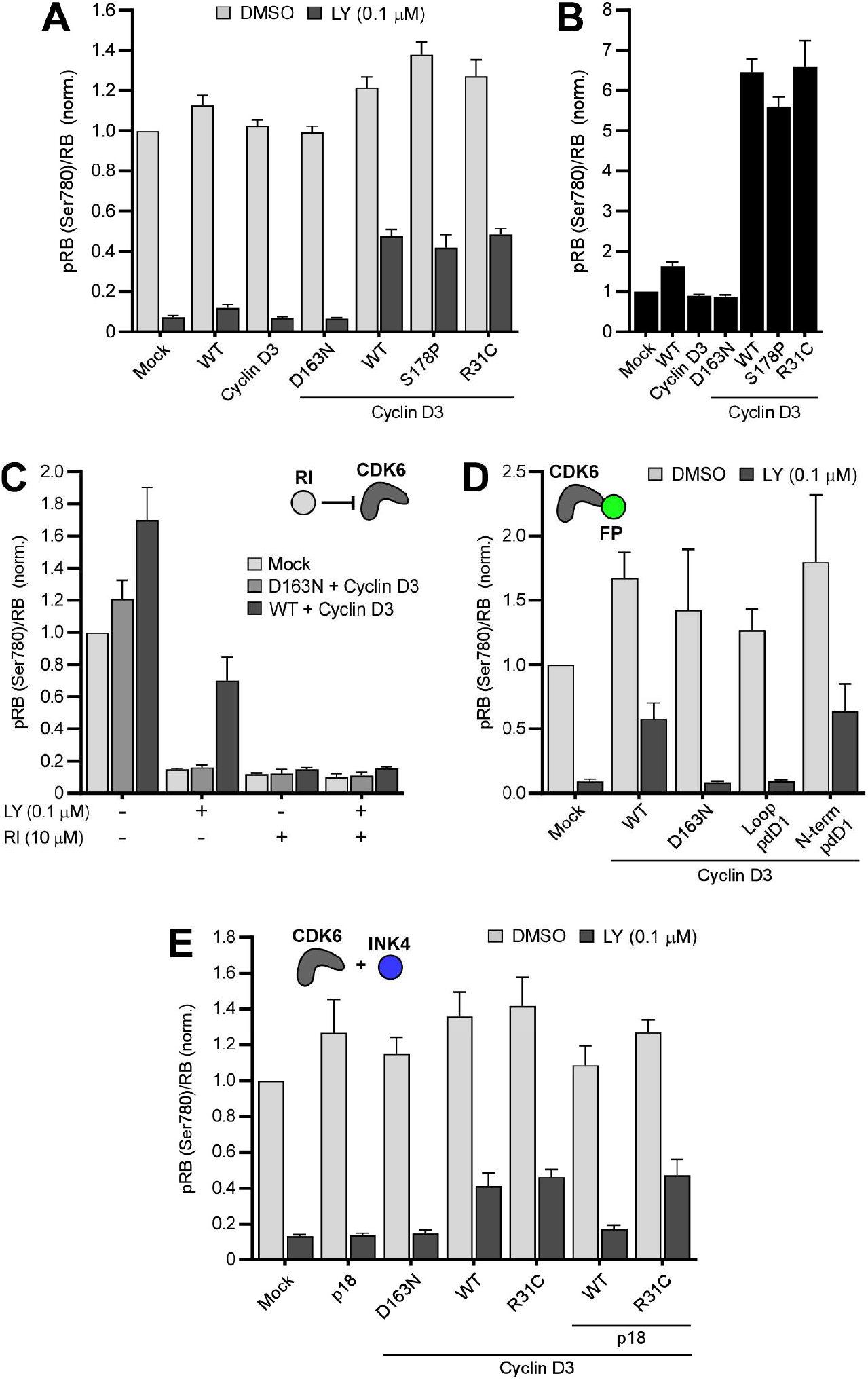
Energy transfer-based assay of CDK6 variants, inhibitors and modulators. (**A**) Ratios of pRB to total RB for MCF-7 cells expressing CDK6 and its variants co-transfected with Cyclin D3. (**B**) LY-treated conditions from (A) normalized to mock LY-treated control. (**C**) Ratios for cells expressing constructs and treated with different combinations of LY and RI. (**D**) Ratios for cells expressing CDK6 containing pdDronpa1 insertions within an internal loop (loop pdD1) or at the N-terminus (N-term pdD1). (**E**) Ratios for cells expressing p18^INK4c^ (p18) in combination with CDK6 or R31C variant and Cyclin D3. (A-E) n=3. Data are mean ± SEM.

Given the efficiency of this method, we applied it to study CDK6 function, inhibition and regulation. We first tested whether LY-treated cells overexpressing CDK6/Cyclin D3 can be used to screen for CDK6 inhibitors. Using RI as a test compound, we found that this is indeed the case (**Fig. 5C**). We also examined genetically-engineered versions of CDK6. Fluorescent proteins are commonly fused to enzymes as reporters of cellular levels/location or as functional regulators (Bajar et al., 2016; Rodriguez et al., 2017; Zhou et al., 2017). These modifications can however be associated with a negative impact on protein function (Patel et al., 2019; Snapp, 2009). We introduced sequences encoding the photoactive pdDronpa1 protein (Zhou *et al*., 2017) at two separate sites of CDK6 to examine if insertion at either site alters activity. pdDronpa1 is a GFP-like protein domain that has been previously used to generate photoswitchable (ps) kinases. Ps-kinases, such as psCDK5, contain pdDronpa1 fused to the N-terminus and inserted in an internal loop (Patel *et al*., 2019; Zhou *et al*., 2017). We introduced pdDronpa1 at the N-terminus of CDK6 as well as within an internal loop (located between helices α_F_ and α_G_, homologous to insertion in psCDK5 (Zhou *et al*., 2017), **Supplementary Fig. S6**) as two separate constructs. We found that the N-terminal insertion was well tolerated; however, insertion into the internal loop site completely abolishes activity (**Fig. 5D**). Lastly, we explored the effects of protein modulators of CDK4/6 function by co-transfecting the INK4 inhibitor protein p18INK4c (p18). INK4 proteins are known to specifically inhibit and modulate CDK4/6 activity by disrupting the cyclin interaction (Nebenfuehr et al., 2020; Parry *et al*., 1999). We transfected p18 alone or in combination with CDK6/Cyclin D3 or CDK6-R31C/Cyclin D3. The R31C mutant was included as it has previously been shown to reduce binding of INK4 proteins to CDK6 (Hu *et al*., 2011). Indeed, we found that p18 was able to effectively inhibit CDK6 but unable to inhibit the R31C mutant (**Fig. 5E**). Overall, these findings establish an assay to test genetically modified CDK6 variants and the effect of CDK6 modulators on kinase activity.

## DISCUSSION

The organization of biological systems into networks permits the emergence of complex functions from a finite number of components and interactions. Kinase-substrate interactions have been previously systematically analyzed towards inference of the global network and targeted perturbation of disease-related network elements. The objectives of our study were reductionist as we focused on local motifs and *in situ* assays. Specifically, we addressed three questions through a combination of bioinformatics and experiments: How common are cKSRs in experimentally validated Ser/Thr and Tyr phosphorylation events? What are the most common motif and family subgroup topologies? And, for the prototypical convergent CDK4/6 kinase pair, how can an individual kinase in the motifs be studied experimentally?

To answer the first two questions, we systematically mapped human phosphorylation sites from the comprehensive PSP. Information in the experimentally validated PSP reflects on a range of factors that govern specificity in the phosphorylation network, such as evolved substrate recognition capabilities or protein localization in subcellular compartments or on adaptor proteins (Creixell et al., 2018; Ubersax and Ferrell, 2007). We found that a large number of human substrates are phosphorylated by more than one kinase. Furthermore, a majority of human kinases participate in cKSRs, and for both Ser/Thr and Tyr kinases a wide variety of interactions were observed. Finally, in many cases the kinases that act on a common substrate are from the same family subgroup and co-expressed in a large fraction of analyzed cell models. This data collectively points to an abundance of convergent phosphorylation interactions that have the potential to drive multi-site phosphorylation events (associated with amplification, crosstalk and feedback) or redundant phosphorylation events (attributed as a source of diversity and robustness). Our analysis is novel and complementary to kinase-substrate enrichment analysis (Casado et al., 2013), which exploits known KSRs to quantify the strength of these interactions in samples under varying experimental conditions. The objective of our work was to identify the basic properties of cKSRs as a basis for further sequence and coregulation analysis (Creixell *et al*., 2018; Humphrey et al., 2013), or for identification of kinases and substrates involved in relevant network motifs, as demonstrated for fundamental reciprocal phosphorylation events. The observation that related kinases phosphorylate substrates has here motivated the development of the experimental strategy that does not rely on kinase-specific inhibitors.

To answer the third question, we studied the convergent CDK4/6 pair under conditions of kinase inhibition. ATP-competitive CDK4/6 inhibitors have led to major improvements in the survival of breast cancer patients. Resistance has emerged in the patient population and, albeit studied intensively, no resistance inducing kinase mutations or substrate protein mutations have been observed. In turn, resistance was attributed to upregulation of CDK6 levels (Cornell *et al*., 2019; Li *et al*., 2022; Li *et al*., 2018; Yang *et al*., 2017). We hypothesized that the interplay of expression level and drug sensitivity can also be explored to design an *in situ* assay for CDK6. We performed these experiments in cells expressing high levels of CDK4 to provide a test bed in which this input entirely masked effects of initial CDK6 overexpression. Ultimately, the generated experimental strategy allowed us to quantify the activity of genetically modified CDK6 kinases, small molecule inhibitors and inhibitory accessory proteins in living cells. The emerging assay complements existing approaches to CDK6 that were in conducted *in vitro* (Bockstaele *et al*., 2009; Li *et al*., 2022; Soamalala *et al*., 2020). It also opens the door to more rapid protein engineering approaches that require quantification of cellular CDK6 activity. For instance, we showed that unlike CDK5 (Zhou *et al*., 2017), CDK6 is not amenable to internal pDronpa1 modification. The ability to perform these experiments with inhibitors from different chemical classes (as demonstrated here for LY and RI) allows separating off-target effects (e.g., from inhibition of upstream kinases). Several potential mechanisms for inhibitor resistance in CDK6-amplified cells have been proposed. In the first mechanism, kinases at elevated protein levels are not efficiently inhibited at tuned drug concentrations, and consequently differential signals are recorded between cells with low (e.g., WT cells) or high CDK6 levels (e.g., CDK6 overexpressing cells). In a second mechanism, resistance of CDK6 to inhibitors may be modulated by accessory proteins (Li *et al*., 2022). Our data suggest that the first mechanism is at play here as we observed a canonical inhibitory function of p18 in the presence of LY leading to reduced pRB levels.

Overall, we here analyzed phosphorylation maps using a convergence-centric approach and, in a prototypical experimental system, dissected convergence. Many diverse cKSRs were observed despite the well documented bias of experimental phosphorylation datasets towards well studied kinases (Buljan et al., 2020; Colinge *et al*., 2014; Invergo and Beltrao, 2018). Our experimental result was achieved even in the absence of kinase-specific inhibitors (e.g., an inhibitor specific for CDK6 over CDK4) or knock-out cell models of the individual converging kinases. Similar limitations may apply to many other scenarios involving convergent kinases. We developed a cellular assay for CDK6 that is desirable as experiments in cells may reflect on a complex natural environment of enzymes and an impact of cellular environment on drug properties. Quite likely redundancy and convergence also exist in other post-translation modification networks, and our work may provide clues as to how to analyze these emerging phenomena.

## ACKNOWLEDGEMENTS

We thank Q. Li and S. Chandarlapaty for advice on experimental design, M. Michael and D. Lynn for advice on databases and expression level analysis. This study was supported by grants of the Australian Research Council (FT200100519 and DP200102093, to H.J.), the National Health and Medical Research Council (APP1187638, to H.J.) and a PhD Top-up Scholarship by the Juvenile Diabetes Research Foundation Australia (to C.G.). The Australian Regenerative Medicine Institute is supported by grants from the State Government of Victoria and the Australian Government. The EMBL Australia Partnership Laboratory (EMBL Australia) is supported by the National Collaborative Research Infrastructure Strategy (NCRIS) of the Australian Government. MicroMon of Monash University provided Sanger sequencing services.

## AUTHOR CONTRIBUTIONS

Conceptualization, C.G. and H.J.; Formal Analysis, R.S. and H.J.; Funding Acquisition, C.G. and H.J.; Methodology, C.G., R.S. and H.J.; Project Administration, H.J.; Investigation, C.G.; Data curation, C.G., R.S. and H.J.; Supervision, H.J.; Visualization, C.G. and H.J.; Writing - Original Draft, C.G. and H.J.; Writing - Review and Editing, H.J.

## Supplementary Information

**Supplementary Tables S1 to S12:** Please see separate Supplementary Data (Excel format).

**Supplementary Table S13:** Reciprocal phosphorylation loops. Loops of the topology shown in Supplementary Fig. S1 were identified in the cKSR data and literature searched for prior mention (see Methods for details).

| Kinase A | Kinase B | Reaction A to B | Reaction B to A | Prior mention |
| --- | --- | --- | --- | --- |
| Abl | MST1 | Tyr | Ser/Thr | N |
| Akt1 | CDK2 | Ser/Thr | Ser/Thr | N |
| Akt1 | CK2A1 | Ser/Thr | Ser/Thr | N |
| Akt1 | GSK3A | Ser/Thr | Ser/Thr | N |
| Akt1 | mTOR | Ser/Thr | Ser/Thr | Y |
| Akt1 | PIKFYE | Ser/Thr | Ser/Thr | N |
| Arg | PDGFRB | Tyr | Tyr | N |
| ASK1 | PDK1 | Ser/Thr | Ser/Thr | Y |
| Btk | ITK | Tyr | Tyr | Y |
| Btk | TEC | Tyr | Tyr | Y |
| CDK1 | CDK7 | Ser/Thr | Ser/Thr | N |
| CDK1 | Chk1 | Ser/Thr | Ser/Thr | Y |
| CDK1 | CK2A1 | Ser/Thr | Ser/Thr | N |
| CDK1 | Wee1 | Ser/Thr | Tyr | Y |
| CDK2 | CDK7 | Ser/Thr | Ser/Thr and Tyr | Y |
| Chk2 | TTK | Ser/Thr | Ser/Thr | N |
| CK2A1 | ERK2 | Ser/Thr | Ser/Thr | N |
| EGFR | ERK1 | Ser/Thr and Tyr | Ser/Thr | Y |
| EGFR | ERK2 | Ser/Thr and Tyr | Ser/Thr | Y |
| ERK1 | Lck | Ser/Thr | Tyr | Y |
| ERK1 | MEK1 | Ser/Thr | Ser/Thr and Tyr | Y |
| ERK2 | MEK1 | Ser/Thr | Ser/Thr and Tyr | Y |
| FAK | Ret | Tyr | Tyr | Y |
| GSK3B | Src | Ser/Thr and Tyr | Tyr | N |
| ITK | TEC | Tyr | Tyr | Y |
| MEK1 | RAF1 | Ser/Thr | Ser/Thr | Y |
| mTOR | p70S6K | Ser/Thr | Ser/Thr | Y |
| PKCD | Src | Ser/Thr | Tyr | Y |

**Supplementary Table S14:**
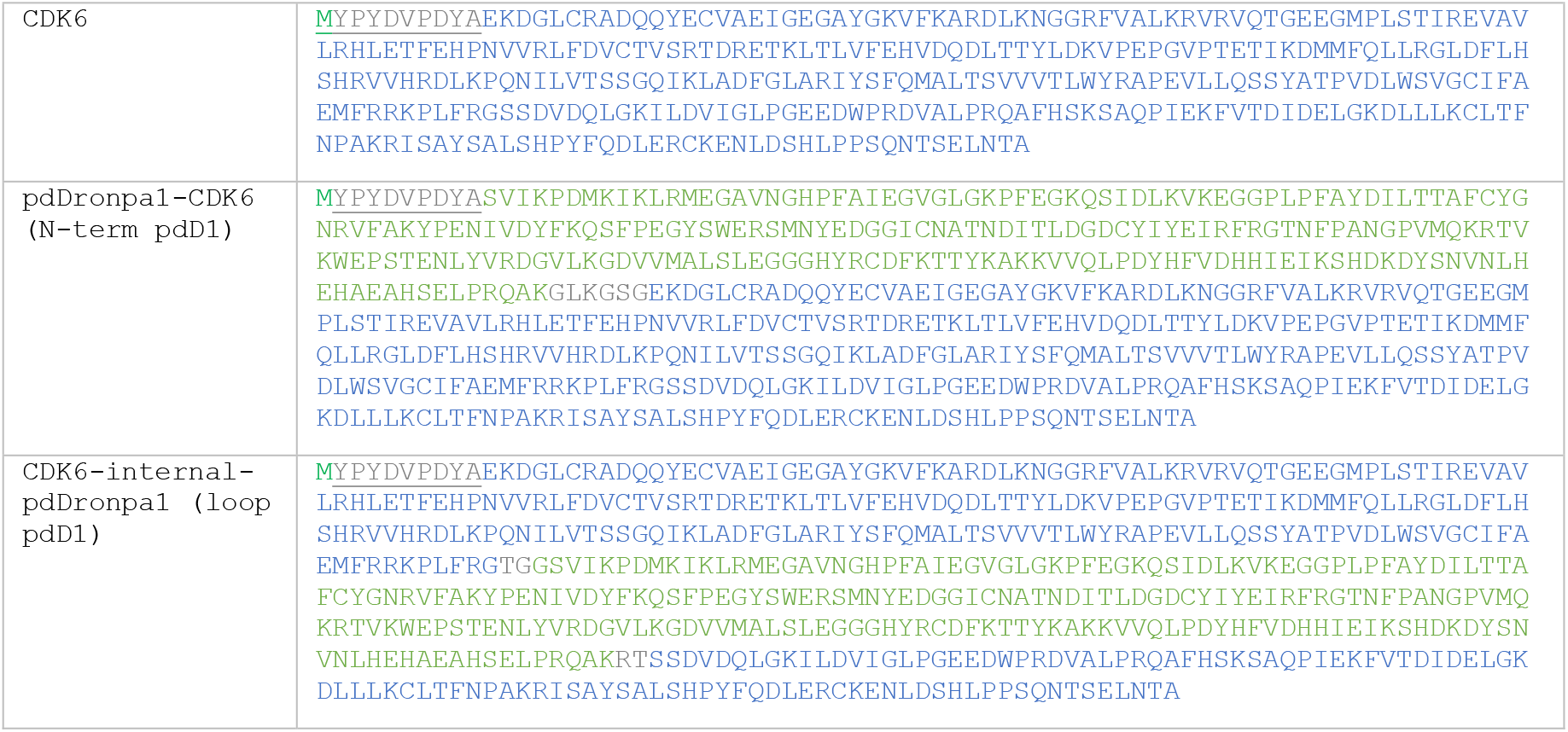
Protein sequences of WT CDK6 and engineered CDK6 variants. Color code: Start codon. <u>HA tag.</u> WT human CDK6. Linkers. pdDronpa1.

**Supplementary Figure S1:**
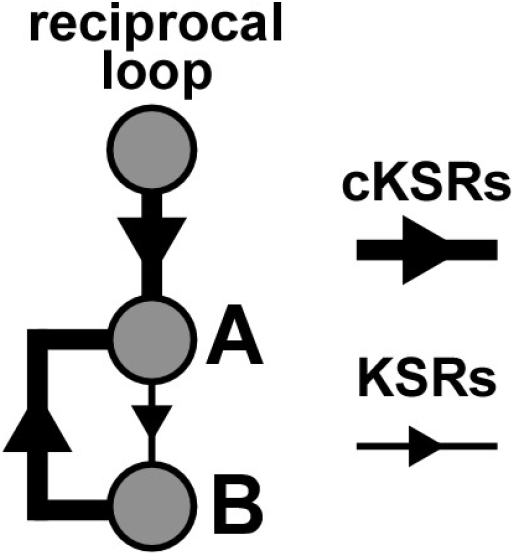
Topology of reciprocal phosphorylation loops. The general cKSR dataset was analyzed to identify all kinases A that receive two inputs and whose substrates B are phosphorylating these kinases reciprocally. The identified kinase pairs are listed in Supplementary Table S13.

**Supplementary Figure S2:**
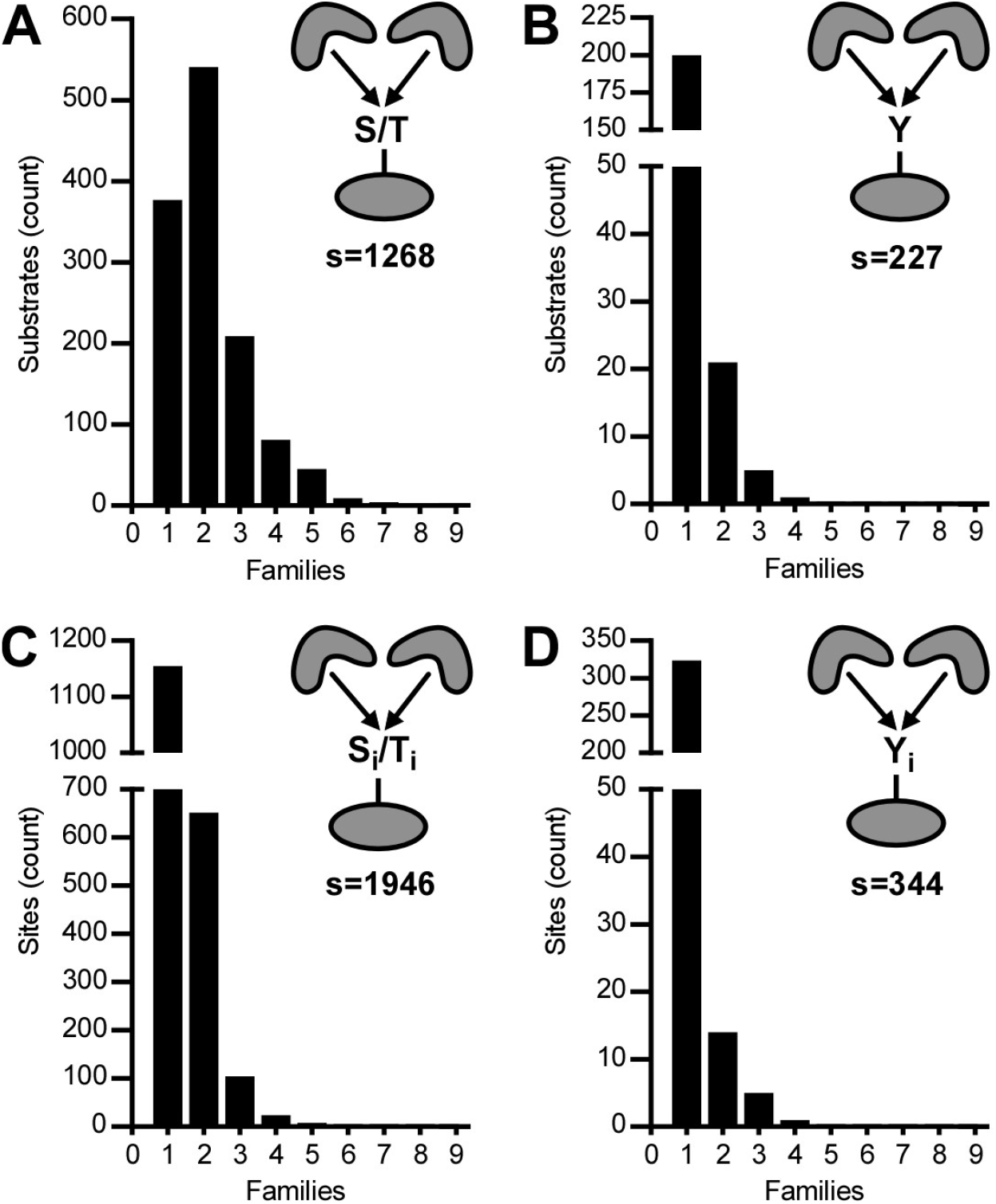
Distribution histograms of kinase families participating in convergent Ser/Thr phosphorylation (A, C) and Tyr phosphorylation (B, D). Bars indicate the number of substrates (A, B) or sites (C, D) that are phosphorylated by members of theindicated number of kinase families. s: Analyzed number of substrates or sites.

**Supplementary Figure S3:**
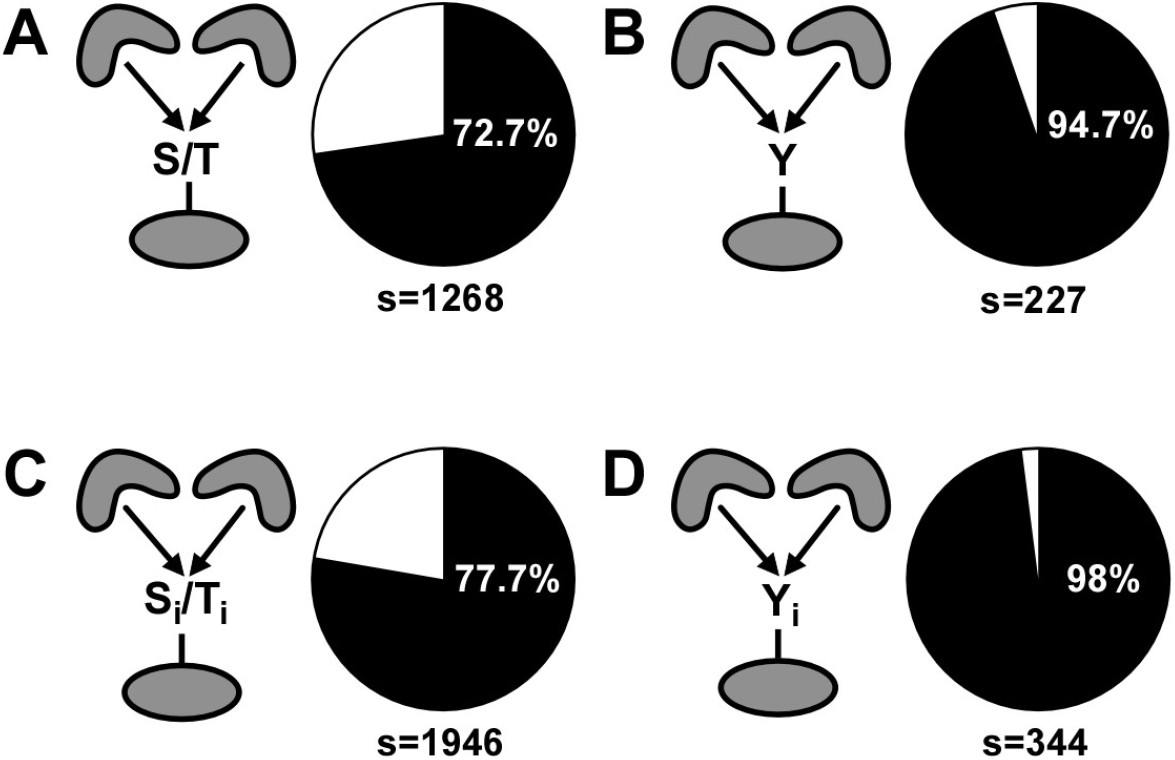
Occurrence of phosphorylation events that include ≥two kinases from the same family subgroup. Pie charts indicating the percentage of substrates that are phosphorylated by ≥two kinases from at least one family subgroup, for general and site-specific Ser/Thr phosphorylation (A, C) and general and site-specific Tyr phosphorylation (B,D). s: Analyzed number of substrates or sites.

**Supplementary Figure S4:**
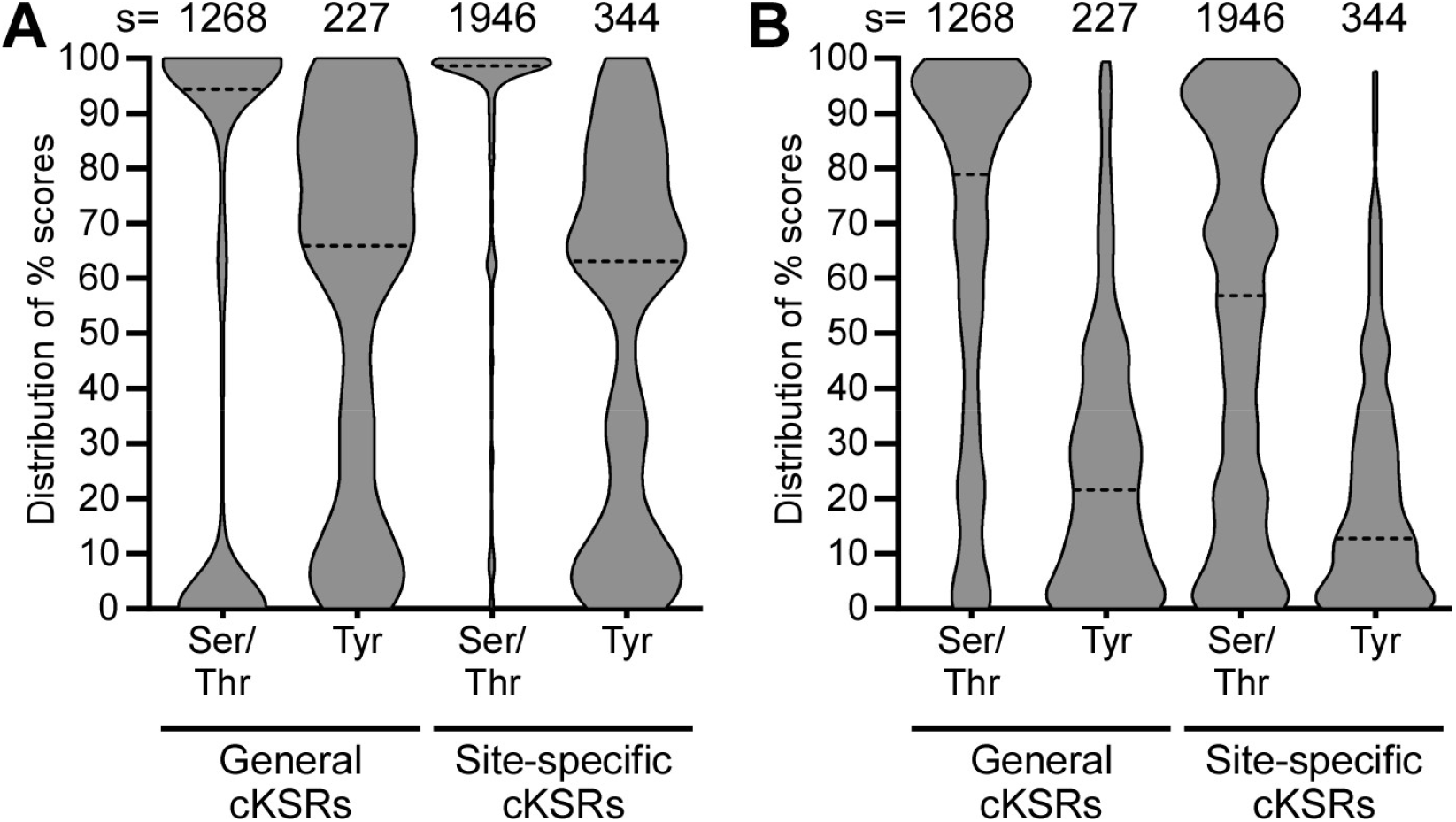
Co-expression analysis at different TPM thresholds. (**A**) TPM threshold of 5. (**B**) TPM threshold of 20. Co-expression scores and median (dashed line) are defined as in Fig. 2. s: Analyzed number of substrates or sites.

**Supplementary Figure S5:**
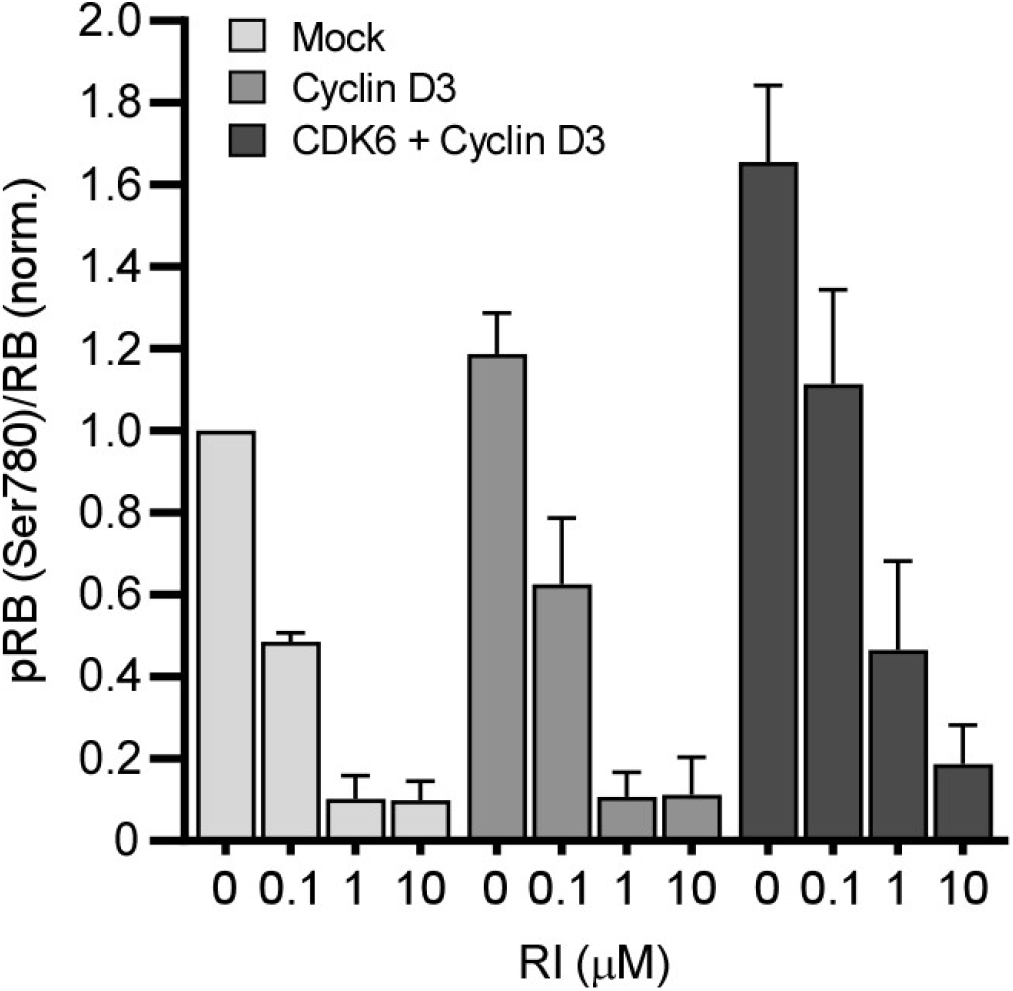
Kinase inhibition with RI reveals CDK6 activity. Energy transfer-based assay to detect ratios of pRB to total RB for MCF-7 cells expressing CDK6 and Cyclin D3 treated with increasing concentrations of RI. n=3. Data are mean ± SEM.

**Supplementary Figure S6:**
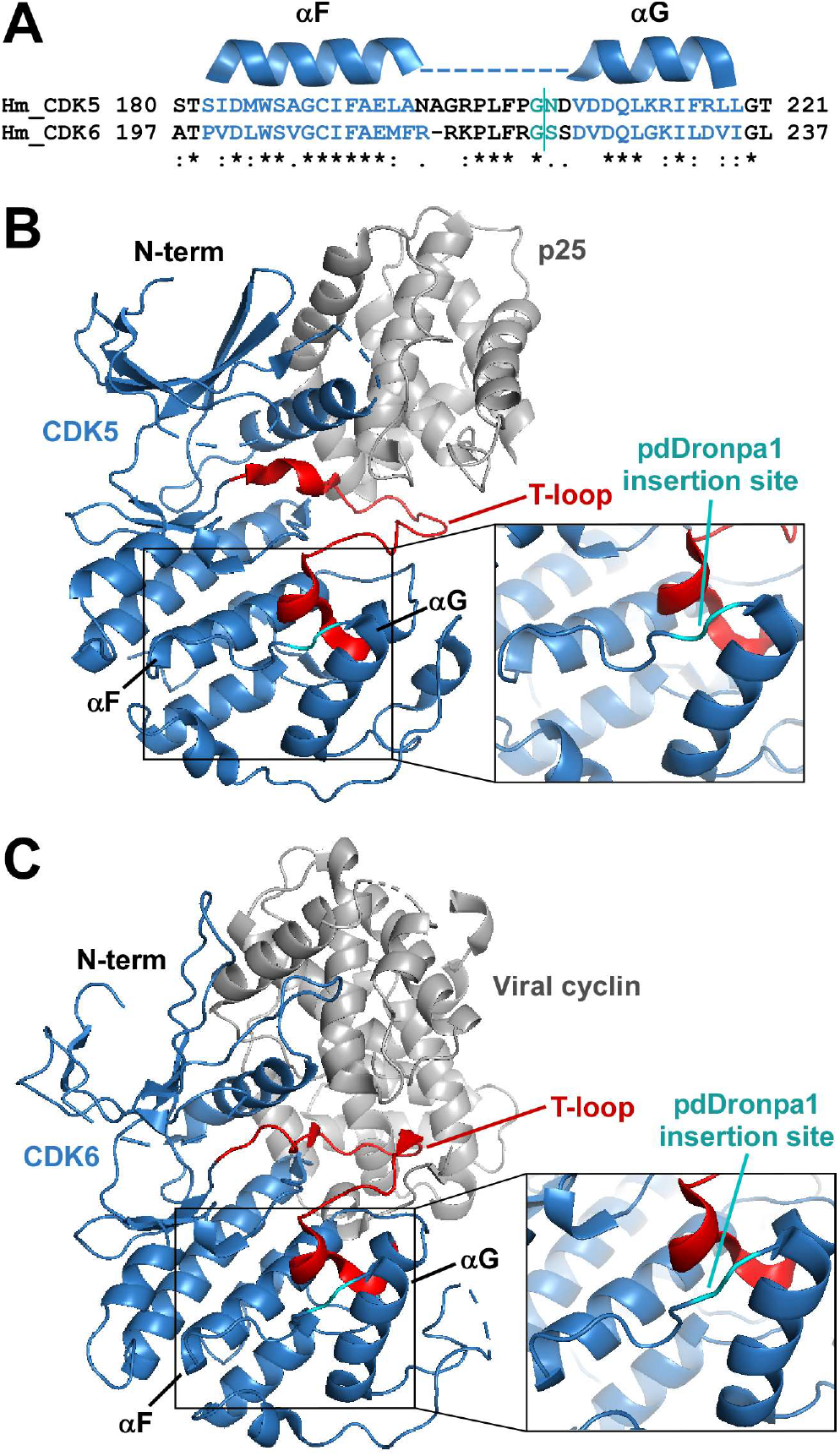
Analogous insertion sites of pDronpa1 in CDK5 and CDK6. (**A**) Protein sequence alignment of human CDK5 and CDK6. pdDronpa1 internal loop insertion site highlighted in cyan (for CDK5, the insertion is equivalent to that of psCDK5 (Zhou *et al*., 2017)). (**B** and **C**) CDK5-p25 and CDK6-viral cyclin complex structures with the insertion sites highlighted (PDB-IDs: 1H4L for CDK5 and 1JOW for CDK6).

